# *Samplify*: A versatile tool for image-based segmentation and annotation of seed abortion phenotypes

**DOI:** 10.1101/2025.09.18.677122

**Authors:** Heinrich Bente, Ronja Lea Jennifer Müller, Andreas Donath, Dirk Walther, Claudia Köhler

## Abstract

Automated seed phenotyping has wide applications in research and agriculture and relies on easy-to-use platforms and pipelines. Seed phenotyping in the model species *Arabidopsis thaliana* poses a significant challenge due to the large number of tiny seeds produced by individual plants, which are difficult to manually separate and count. Manual counting methods are time-consuming and prone to user bias, particularly for subtle phenotypic changes. To address these limitations, we developed *Samplify*, a scalable, automated pipeline for seed segmentation and classification. By integrating classical image processing techniques with Meta’s Segment Anything Model (SAM), *Samplify* effectively segments *Arabidopsis* seeds, even in dense clusters where conventional methods fail. To demonstrate its versatility, we quantified the seed abortion occurring in interploidy crossings in Arabidopsis, often referred to as ‘triploid block’. *Samplify* includes a Random Forest classifier trained on a set of computed seed shape features that enables the categorization of seeds into normal, partially aborted, and fully aborted seeds, automating the manual classification process. The tool, designed as a command-line application, significantly reduces manual annotation workload. Our validation across multiple datasets demonstrates high segmentation and classification reliability, making *Samplify* a valuable resource for the plant research community.

## Introduction

Seed development in flowering plants (angiosperms) is initiated by a process known as double fertilization, during which two haploid sperm cells from the pollen fertilize the two female gametes, the egg cell and the central cell, resulting in the formation of the embryo and endosperm, respectively (Dresselhaus et al., 2016). In angiosperms, the central cell is predominantly homodiploid, forming a triploid (3x) endosperm that inherits two copies of the maternal and one of the paternal genome (2m:1p). This specific genome ratio in the endosperm is essential for the survival of the embryo in the majority of angiosperms. In *Arabidopsis thaliana*, as in most angiosperms, the endosperm initially develops as a coenocyte for a defined number of mitotic cycles, after which it starts to cellularize (Boisnard-Lorig et al., 2001). Endosperm cellularization is critical and ensures proper embryo growth; cellularization failure will cause embryo arrest and seed abortion (Hehenberger et al., 2012). Changing the parental genome ratio in the endosperm affects endosperm cellularization and embryo development in F1 seeds. Pollinating a tetraploid maternal plant with a diploid pollen donor results in 3x embryos and 5x endosperms. The endosperm of those seeds cellularizes precociously, leading to smaller seeds that in Arabidopsis are nevertheless viable (Scott et al., 1998). The reciprocal cross of maternal diploid plants with pollen from a tetraploid plant results in 3x embryos and 4x endosperms. These seeds are characterized by delayed or failed endosperm cellularization, resulting in seed abortion. This phenomenon is often referred to as ‘triploid block’ (Marks, 1966, Köhler et al., 2010) and acts as a postzygotic hybridization barrier in a wide range of taxa (Bente and Köhler, 2024, Coughlan, 2023). Thus far, the genetic basis of the triploid block has been mainly investigated in Arabidopsis (Kradolfer et al., 2013, Martinez et al., 2018, Satyaki and Gehring, 2019, Batista et al., 2019).

Quantifying seed abortion typically involves manually photographing seed populations and categorizing seeds based on deviations in shape and color. Abnormal seeds are classified as either partially or fully aborted: partially aborted seeds appear misshapen but contain a developed embryo that often germinates, while fully aborted seeds are dark brown and shriveled and fail to germinate. However, this manual assessment is time-consuming and inherently subjective, introducing variability and potential biases into the data.

To minimize observer bias, we developed an analysis pipeline based on automated image analysis, a critical step in quantifying seed abortion. Image analysis generally can be divided into two separate steps: segmentation, the partitioning of a given image into meaningful regions, and classification of the afore-identified regions. While classification involves relatively simple tasks such as size measurements, segmentation is the crucial prerequisite. Image segmentation itself can be typically classified into semantic segmentation, in which each pixel of an image gets a class label, instance segmentation, which identifies objects of the same class, whereas panoptic segmentation relies on a combination of the two (Kirillov, 2019). Segmenting images showing plant seeds is challenging because of high numbers and densely clustered seeds of different shapes that occasionally overlap. To circumvent this problem, different approaches were previously applied, ranging from manually dispersing seeds prior to imaging (Benjamaporn and Chomtip, 2017, Kurtulmuş, 2021), to the application of large particle flow cytometry for sorting single seeds (Morales et al., 2020), or using robotic approaches combining single seed sorting and imaging at the same time (Jahnke et al., 2016, Krzyszton et al., 2024, Klasen et al., 2025). Nonetheless, purely image-based methods have become more prominent, as they offer affordable and standardized analysis without requiring specialized equipment. These methods have been mainly used for larger plant seeds, such as sugar beet (Wang et al., 2023) and soybean (Wei et al., 2023).

Alongside traditional image analysis tools, a range of specialized seed phenotyping pipelines have been developed, primarily targeting cereal crops and other large-seeded species. Notable examples include ‘SmartGrain’ (Tanabata et al., 2012) and ‘GrainScan’ (Whan et al., 2014), which have been successfully applied to species such as wheat, rice, millet, and *Brachypodium*. These tools are often optimized for high-resolution input from flatbed scanners, which provide consistent lighting and minimize shadows, allowing for relatively straightforward segmentation methods such as global or adaptive thresholding. However, such approaches typically assume well-separated seeds and tend to remove or misclassify dense clusters of touching or overlapping seeds, which limits their applicability to species where seeds are small and more prone to aggregation during imaging. While handheld scanners have been explored as a more flexible imaging option, particularly for legumes like beans and peanuts (Huang et al., 2024), these devices often lack the resolution required to reliably capture the fine morphological features of smaller seeds. Additionally, many of these pipelines produce only basic shape or color descriptors (e.g., area, circularity, RGB values), placing the burden of biological interpretation and downstream statistical analysis on the user. More recently, the integration of artificial intelligence (AI) and deep learning has advanced seed image analysis by improving segmentation accuracy and enabling feature extraction from more complex or noisy images. These methods offer the potential to overcome challenges such as overlapping seeds and non-uniform lighting conditions. However, most AI-driven applications have so far been focused on larger-seeded crops like rice, maize, or medicinal plants such as *Scutellaria baicalensis* (Jinfeng et al., 2023, Keling et al., 2023) with relatively little attention given to small-seeded species.

Thus, in the case of Arabidopsis seeds, which are usually small and come in high numbers, previous studies used manual quantification of phenotypes based on pictures, or dispersing seeds before imaging (Merieux et al., 2021, Ren et al., 2019, Herridge et al., 2011).To ease quantification of seed phenotypes, we developed *Samplify*, a command line tool designed for automated segmentation and classification of Arabidopsis seeds. The name reflects the integration of Meta’s ‘Segment Anything Model’ (<u>SAM</u>), and the purpose of sim<u>plify</u>ing otherwise time-consuming and user-dependent seed classification. *Samplify’s* segmentation pipeline is designed in a hybrid-segmentation approach, balancing computational efficiency and segmentation accuracy by combining classical image processing techniques with advanced transformer-based models. The segmentation of regions where seeds are sparsely distributed and do not touch each other is achieved via traditional processing methods, namely Otsu thresholding(Otsu, 1979). However, in regions with dense clusters of seeds that touch or overlap, *Samplify* utilizes SAM (Kirillov, 2023), a transformer-based model designed for general-purpose segmentation. By leveraging SAM’s capabilities, *Samplify* achieves high segmentation accuracy in challenging regions where conventional methods would typically require manual correction. Previous studies have demonstrated that implementing SAM-based segmentation enhances performance, specifically in medical-imaging (Li et al., 2025, Ma et al., 2024, Wei et al., 2023, Junde et al., 2025). Combining traditional image segmentation with SAM offers a scalable, efficient, and robust segmentation strategy. From each segmented entity, the tool collects multiple features, including shape, color and texture that the user can access for downstream analysis. *Samplify* was specifically designed and tested for the quantification of seed abortion in *Arabidopsis thaliana*, thus we included a Random Forest (RF) classifier that subsequently labels segmented seeds into normal, partially or fully aborted seeds. The RF-model was trained on more than 13000 seeds across 99 images that were all manually annotated, resulting in a robust RF-model for seed abortion prediction. We tested *Samplify* using different independently generated datasets, demonstrating its high accuracy in segmentation and reliable seed abortion prediction. We believe that in combining automated segmentation together with a robust seed abortion RF classifier, *Samplify* reduced user-bias inflicted inaccuracies as well as time-consuming manual counting of such phenotypes, thus providing a valuable tool for the plant community.

## Results

*Samplify* is a purely image-based seed phenotyping platform, requiring users to provide only pre-annotated pictures for training and seed abortion prediction (Fig. 1A). The *Samplify* pipeline consists of two steps, namely a “training”- and “prediction” step, both necessary to achieve automated seed phenotyping. As *Samplify* relies on a machine learning approach, users first need to use their pictures to train a random forest model before being able to start the prediction. The training step consists of multiple smaller steps as indicated by the blue arrows in Fig. 1B.

**Figure 1:**
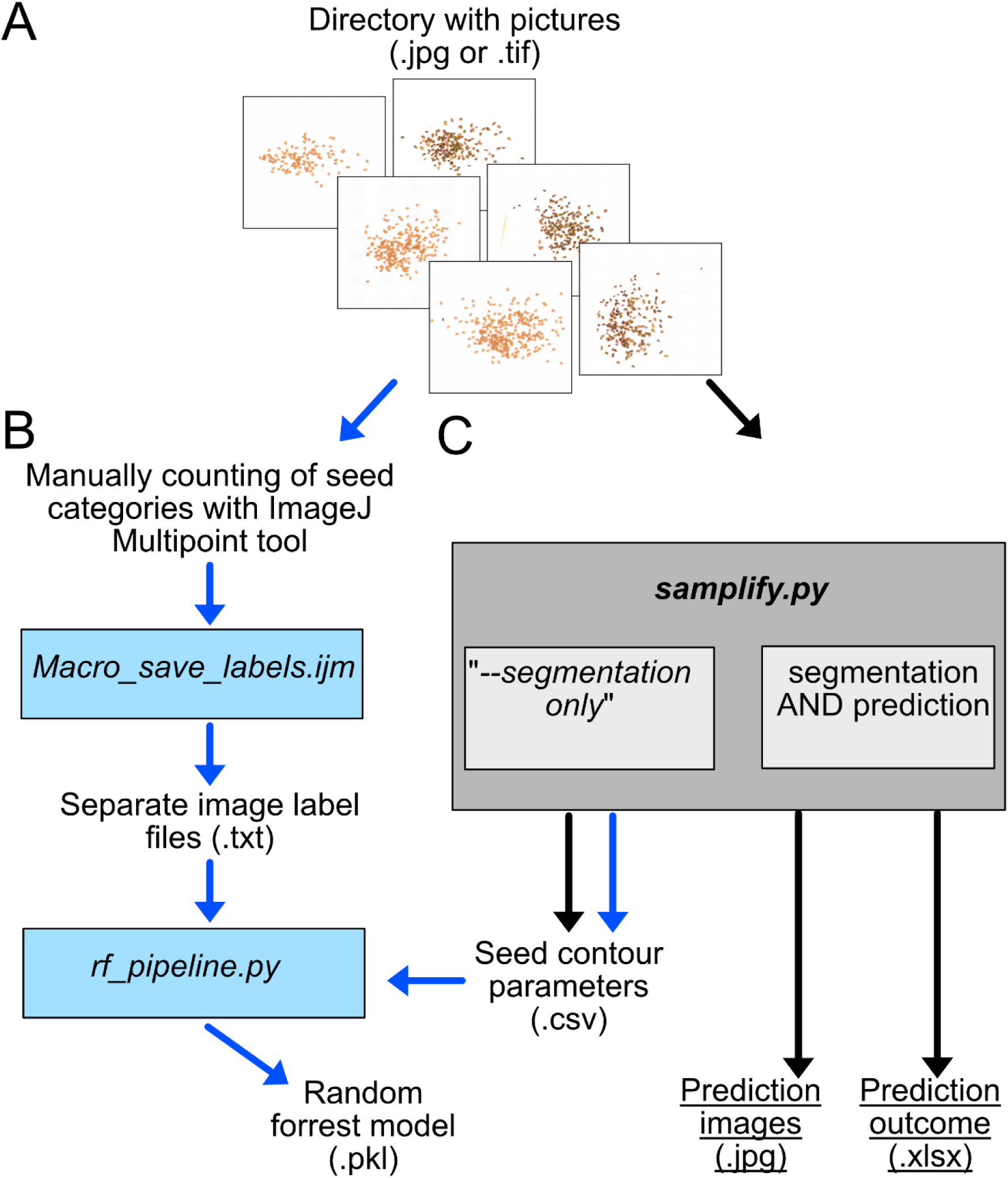
Workflow for *Samplify*-based phenotyping of seed abortion. In order to use Samplify, users have to provide a directory with images in .jpg or .tif format for either training a random forest model or for predicting seed phenotypes. A) For the generation of a random forest model, each picture had to be separately counted using ImageJ multipoint tool as described and the label information has to be extracted using the *Macro_save_labels.ijm* macro of ImageJ. In the same directory, users can run samplify.py --segmentation-only to generate seed contour parameters. The Image label files together with the seed contour parameter file is used to run the python script rf_pipeline.py to generate and test a random forest model (.pkl) including testing parameters of the same. B) By using a prior created random forest model, users can start predicting seed abortion phenotypes on either the same pictures as used for the random forest model for validation or on an independent set of pictures. By default, samplify.py performs a segmentation that outputs a seedparameter.csv file and a prediction outcome in .xlsx format. Moreover, for each input picture, an annotated prediction picture is created in a subdirectory to allow fast validation. Blue arrows indicate the training path, black arrows the segmentation and prediction path. Blue and grey boxes indicate different scripts for each of the tasks utilized, plain text describes different output files, with the file ending in brackets.

Pictures used for training need to be manually annotated, which can easily be done using the Multipoint-tool in *ImageJ* (Schindelin et al., 2012). For annotating pictures for the triploid block, we counted each picture of the training set using the Point-tool, assigning “Counter 0” for “normal seeds”, “Counter 1” for “partially-aborted seeds” and “Counter 2” for “fully-aborted seeds”. To extract the label information of each picture, we used the *ImageJ* Macro *Macro_save_labels.ijm*, which needs to be downloaded and installed in the ImageJ software. The Macro accesses the directory with all annotated TIFF files and creates a TXT-directory containing for each annotated TIFF file a separate image label file (.txt) with coordinates and categories for a given picture. As we opted to train the random forest model only on proper segmented pictures, we added a segmentation-only option in *Samplify*’s main program. Running *samplify.py --segmentation-only* on a directory with training images (either raw JPGs or annotated TIFFs), results in *seed_parameters.csv* containing a table with all segmentation information (Supplementary Table S1) from all images in the input directory.

In the segmentation process, *Samplify* will take each single picture and apply a hybrid segmentation model, trying to identify clusters of seeds and applying Metas “Segment Anything” (SAM) image analysis to identify single seeds. Only for seeds not appearing in a cluster as they are too dispersed, *Samplify* utilizes traditional thresholding techniques to achieve segmentation of all individuals in one picture.

In the final training step, the *rf_pipline.py* file uses the *seed_parameters.csv* file from the *Samplify* segmentation and the TXT-directory of the Separate image label files to create a Random Forest model (.pkl) and testing statistics of that model. The whole training workflow is depicted by blue arrows in Fig. 1B. After users have approved the results of the test model, it can be used for different predictions. This only requires the raw pictures, which should ideally be taken using the same settings as done for training dataset. Users can run samplify.py in a directory containing the pictures to be analyzed. *Samplify* will automatically use the default random forest model *TripBlockDefault_RF.pkl* as long as no other model is specified. As output, *Samplify* will generate a subdirectory called “out” containing the *predicted_images* directory with all *Samplify*-annotated images, as well as two files, a *seed_parameter.csv* file with all picture segmentation and image information. It will furthermore generate a *seed_summary.xlsx* file, which contains the number of annotated total seeds and seeds per phenotype category (“normal”, “partially”, “fully aborted”), as well the relative values in percentage per picture. Moreover, the *seed_summary.xlsx* file contains further information on processing time and prediction confidence.

As *Samplify* was specifically developed for the segmentation and subsequent quantification of seed abortion, we trained an RF model on a comprehensive dataset consisting of 99 pictures, obtained from multiple triploid block experiments, in total containing 13395 single seeds. The distribution across the different seed abortion categories showed that 36.4% of these seeds were manually labeled as normal, 28.5% as partially-aborted and 35.1% as fully aborted seeds (Fig. 2A). A principal component analysis (PCA) across the different manually-annotated seed abortion categories showed that 35.2% and 21% of variation can be dissolved by PC1 and PC2, respectively (Fig. 2B). Furthermore, the PCA revealed that partially aborted seeds were not clearly separated from viable and fully aborted seeds, reflecting these categories are sometimes not clearly distinguishable (Fig. 2B). Training the RF classifier on that dataset resulted in an accuracy of 87.8% across 13064 single individuals that could be segmented properly and assigned to manually-annotated labels. There was only a small percentage of seeds where predicted labels deviated from the actual labels, especially for partially aborted seeds (Fig. 2C). Lastly, we tested the relative feature importance after training the RF classifier. Plotting the top ten most important features to distinguish different seed abortion categories revealed that the overlap ratio, measuring the intersection-over-union (IoU) of the seed shape to an ellipse fitted to its contour, was the top contributor with 15.4%, followed by red and green color parameters. Thus, the RF uses the same features for classification as subjectively assessed by human annotators (Fig. 2D; Supplementary Fig. S2). In conclusion, our trained RF model, based on 13169 manually labeled seeds, demonstrates high accuracy in predicting normal and fully aborted seeds. However, partially aborted seeds are sometimes incorrectly annotated due to their shared features with both normal and aborted seeds.

**Figure 2:**
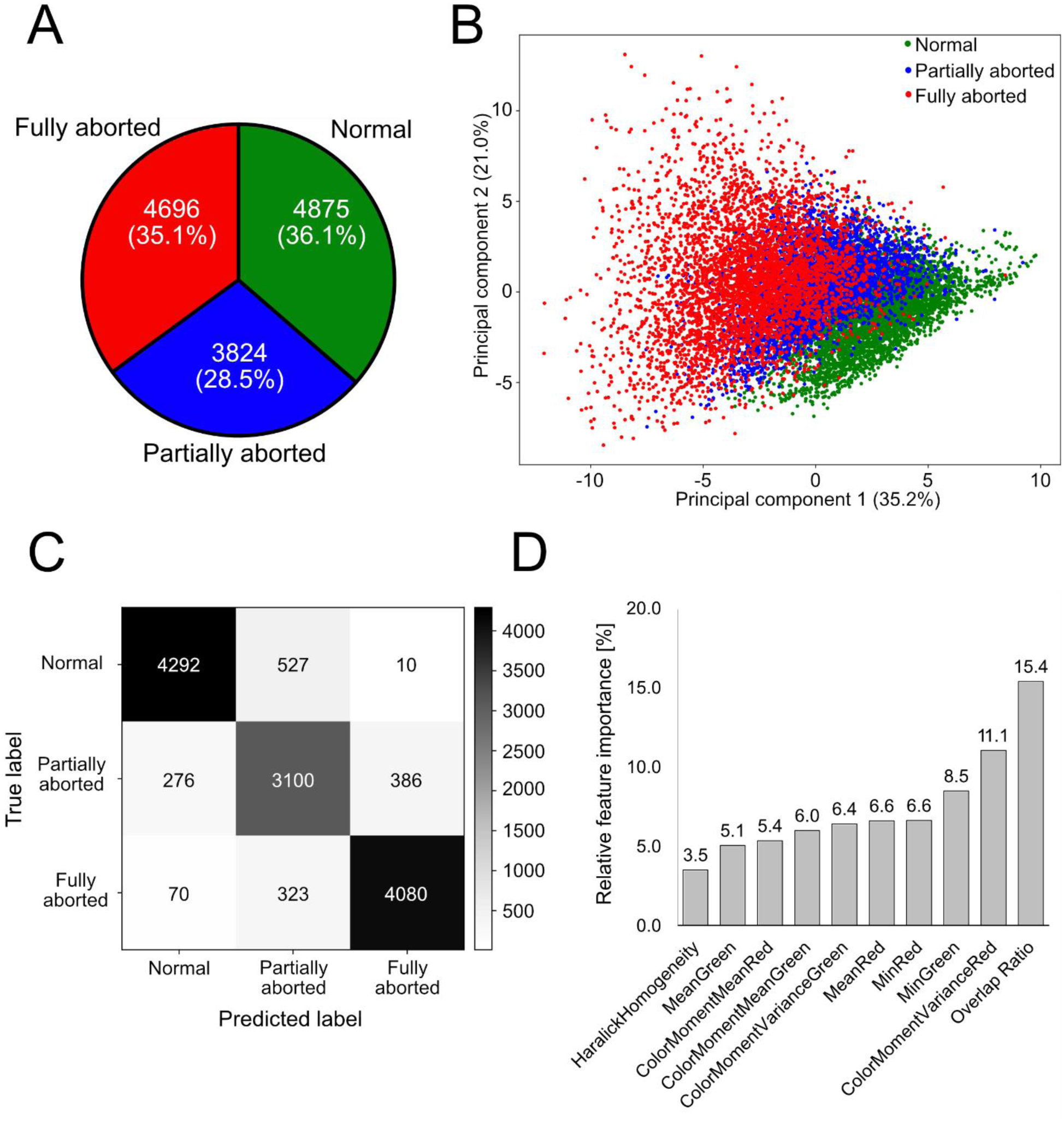
Default trainings dataset for *Samplify*. A) PieChart depicting the total and relative amount of manually annotated seed data from 99 pictures containing a total of 13395 seeds, which contain 42.2% normal, 25.1% partially aborted, and 32.7% fully aborted seeds. B) Principal component analysis (PCA) across different manually annotated seed abortion categories in the training’s dataset. C) Confusion Matrix of 13064 seeds with overlapping label and segmentation info, illustrating the overlap of manually labeled and predicted seeds by *Samplify*. D) Relative feature importance of the top ten parameters used in the random forest (RF) classifier for the default model. A full list of feature importance can be found in Supplemental Fig. S2.

After having successfully established training and validation with the training dataset, we tested *Samplify’s* segmentation and prediction with independent triploid block data. For this test, we included pictures of F1 seeds obtained by crossing Col-0 × Col-0, resulting in normal-looking, light brown, plumb seeds referred to as ‘diploid seeds’. Moreover, we included the same number of pictures from F1 seeds of Col-0 × 4x Col-0, a cross resulting in a high seed abortion rate, referred to as ‘triploid block’ (Scott et al., 1998). These aborted seeds have a variable size and shape and are darker in color, specified as ‘triploid seeds’ (Fig. 3A). The majority of seeds from F1 Col-0 × Col-0 were predicted by *Samplify* as normal seeds, whereas F1 Col-0 × 4x Col-0 were predominantly predicted as partially or fully aborted seeds (Fig. 3B). To obtain quantitative data, we applied *Samplify* using our default random forest model on six replicates each for F1 Col-0 × Col-0 seeds and F1 Col-0 × 4x Col-0 and plotted the relative amount of normal, partially aborted and fully aborted seeds (Fig. 3C). As expected, diploid seeds were predicted to be predominantly (98.24%) normal, in contrast to only 4.3% of triploid seeds. Triploid seeds were predicted to be either partially aborted (20.4%) or fully aborted (75.3%), while these categories were rarely predicted for diploid seeds (<1%, Fig. 3C; Supplementary Table S2). These values are well in line with manual classification of seed abortion based on the same pictures, resulting on average in 99.3% normal, 0.4% partially and 0.3% fully aborted diploid seeds, compared to 1.7% normal, 11.5% partially and 86.8% fully aborted triploid seeds (Fig. 3D; Supplementary Table S3).

**Figure 3:**
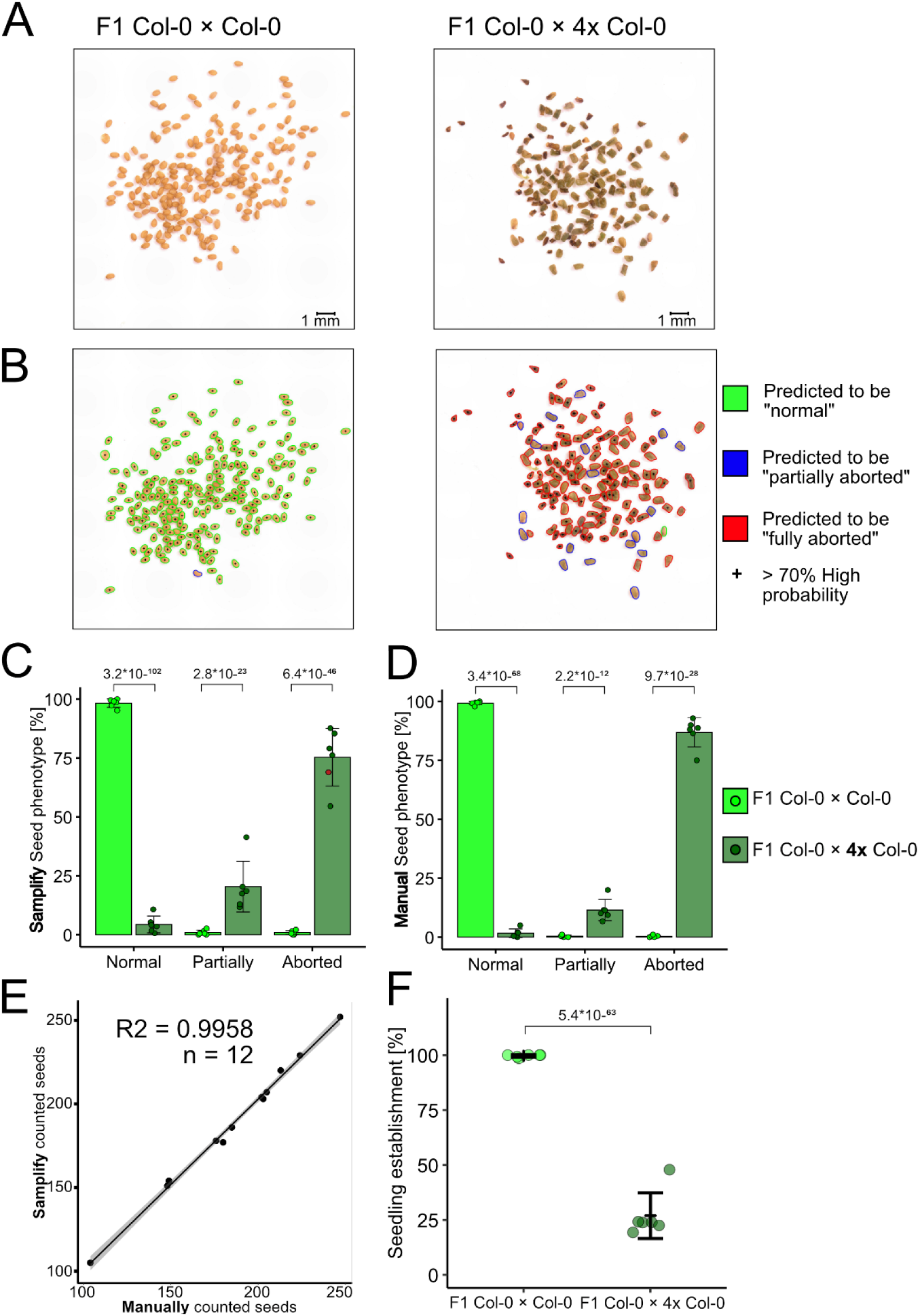
Samplify faithfully segments, annotates, and predicts seed abortion in control crosses. A) Input pictures of F1 Col-0 × Col-0 (left side) and F1 Col-0 × 4x Col-0 (right side) seed populations and its annotation by *Samplify* (B). Relative seed phenotypes from F1 Col-0 × Col-0 and F1 Col-0 × 4x Col-0 annotated by *Samplify* (C) and manually annotated (D). Statistical comparison was done per category by applying generalized linear models (GLMs) with a Poisson distribution, p-values of comparisons are plotted. E) Scatterplot of *Samplify* segmentation against the total number of manually counted seeds per picture. F) Seedling establishment at 4 days of growth in F1 Col-0 × Col-0 and F1 Col-0 × 4x Col-0. Statistical comparison was done using a generalized linear model (GLM) with a Poisson distribution, p-value as indicated. A full list of test statistics and values used can be in Supplementary Tables S2-S5.

To evaluate *Samplify’s* segmentation capability, thus recognizing single seeds in given pictures of seed populations, we compared the total number of manually counted seeds from all pictures *versus* the number of seeds segmented by *Samplify*. Single seeds counted manually and detected by *Samplify* nearly perfectly aligned, resulting in an R^2^ value of 0.9958 across 12 independent seed populations (Fig. 3E). Manual counting resulted in 2236 seeds, whereas *Samplify* identified 2266 seeds, a difference of only 30 seeds and less than 2% variation (Supplementary Table S4).

Finally, we tested whether seeds predicted as normal or partially aborted can germinate, thus correlating the prediction of seed viability with the potential to germinate. Similar to the high proportion of normal diploid seeds predicted as well as manually counted, these seeds had a high germinability of 99.6% over all replicates. In contrast, triploid seeds had, on average, a germination rate of 26.9%, corresponding well to the predicted number of 20.4% partially aborted seeds (Fig. 3F; Supplementary Table S5), which were previously shown to largely keep their potential to germinate (Hehenberger et al., 2012, Batista et al., 2019). Indeed, the prediction of partially aborted seeds based on *Samplify* comes closer to the actual number of germinating seeds than the number based on manual counting (11.5%), demonstrating the high prediction value of *Samplify*. Taken together, these results demonstrate that the automated segmentation and annotation of seed abortion is highly reliable and aligns and even outperforms traditional methods of manual counting, thus providing a fast and less biased way of quantifying seed abortion.

To further test the prediction confidence of *Samplify* across different biological samples, we evaluated its reproducibility by comparing predictions of seed abortion in the same population using different images. We selected a seed population with normal, partially, and fully aborted seeds and imaged these seeds using the same settings; however, after each image, the seeds were newly dispersed for the next image, resulting in multiple pictures of the same seed population. The predicted number of seeds as well as predicted categories was highly similar among the replicates, revealing low technical variation due to different seed positions (Fig. 4A; Supplementary Table S6). To further test the technical variation of *Samplify’s* seed prediction, we measured the per-category reproducibility across six random seed populations, each imaged in seven iterations. Reproducibility was calculated as 1-coefficient of variation (CV), in which CV is the standard deviation (SD) divided by the mean, SD/mean. Reproducibility was assessed for total seed number as well as for each seed phenotype category independently. To account for low numbers, reproducibility was calculated only for those populations that showed an average count >10 in each category. This resulted in an average reproducibility of 0.983 for total seeds (n=6), 0.955 (n=4) for normal seeds, 0.853 (n=3) for partially aborted seeds, and 0.958 (n=4) for fully aborted seeds (Fig. 4B; Supplementary Table S7). Together, this analysis shows high reproducibility for the prediction of total seed number, high reproducibility for normal and fully aborted seeds, and good reproducibility for partially aborted seeds. The fact that partially aborted seeds show less reproducibility compared to the other categories is a consequence of their intermediate phenotype, sharing characteristics of normal and aborted seeds.

**Figure 4:**
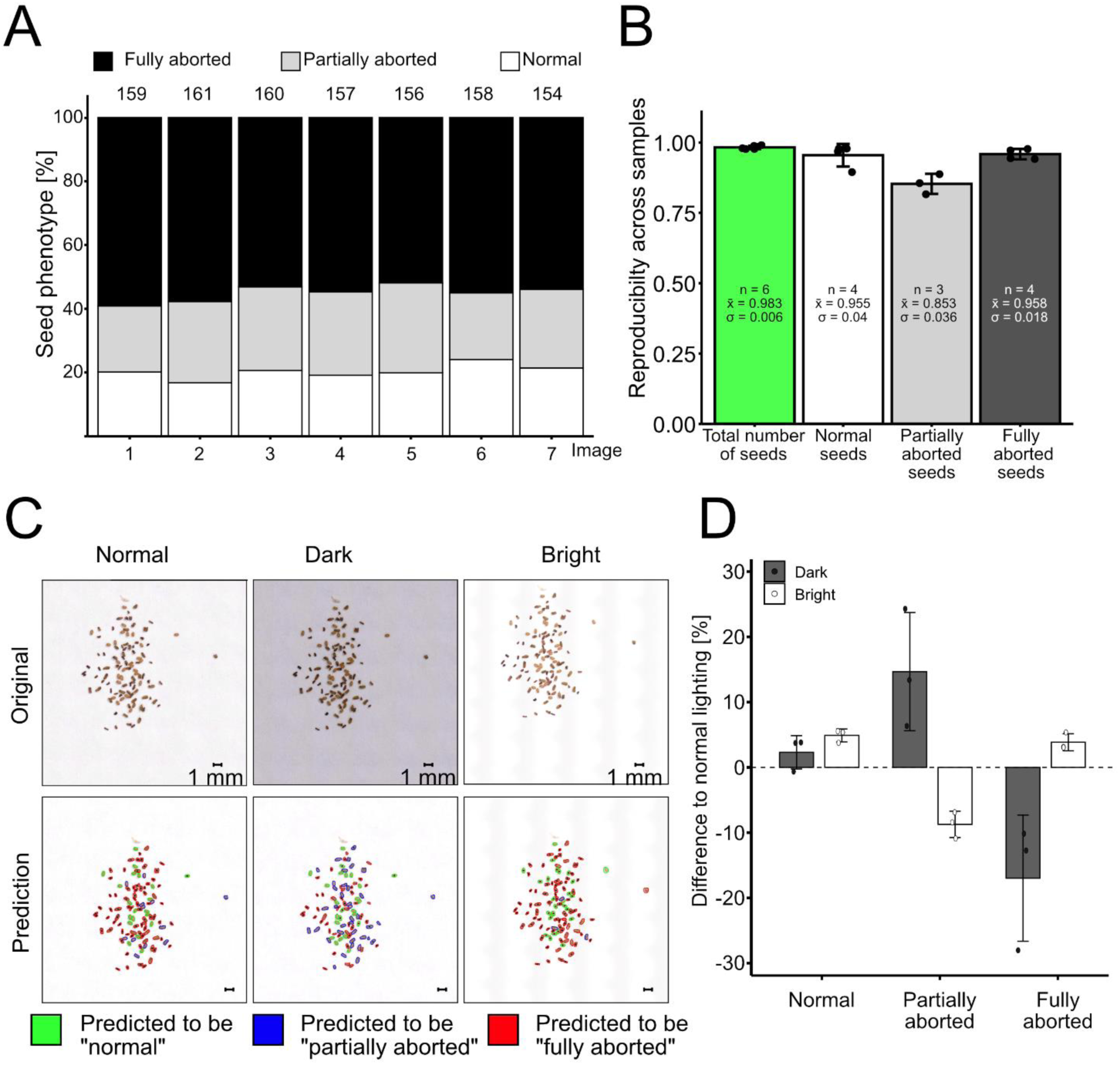
*Samplify* predictions are reproducible across different iterations. A) *Samplify* quantified seed abortion on seven pictures of one seed population. Each picture was taken with the same setting and seeds were randomized in between acquisitions. Numbers on top represent the total amount of seeds detected by *Samplify*. B) Reproducibly across different seed populations per scored category. Reproducibility was calculated by 1-coefficient of variation (CV). Whereby CV is the standard deviation (σ) divided by the mean (x̄) of each seed population in seven different pictures. To account for low numbers, reproducibility was plotted only for those populations that showed an average count >10 in each category, n represents the number of populations matching this criterion. A full table of raw counts and reproducibility calculations can be found in Supplemental Table S6-S7. C) Original pictures (upper panel) and *Samplify* prediction pictures (lower panel) of the same seed population at normal lighting conditions, under-exposed settings ("Dark"), and over-exposed setting ("Bright"). D) Change in relative seed phenotype predictions at different lightning conditions as shown in (C). Three seed populations were imaged at three different light conditions and the relative change in seed phenotype per category is shown. Error bars represent standard deviation. Relative values are denoted in Supplementary Table S8.

We further tested whether reproducibility would be affected by imaging conditions, in particular, different light exposure and resolution. We tested Samplify’s seed phenotype prediction using different lighting conditions, by comparing our normal lighting condition with under-exposed images referred to as “dark” and over-exposed conditions called “bright” (Fig. 4C). *Samplify* does a background normalization, thus all images were adjusted, resulting in perfect white backgrounds in the prediction images (Fig. 4C, lower panel). The fraction of normal seeds was the least affected category, with on average 2.3% more normal seeds predicted in dark conditions and 4.9% more in bright conditions. The prediction of partially aborted seeds was more strongly affected by lighting conditions, with an average increase of 14.7% in dark conditions and a decrease of 8.7% in bright conditions. This substantial change in partially aborted seed prediction was accompanied by a corresponding reduction in the annotation of fully aborted seeds, which decreased by 17% in dark conditions and increased by 3.9% in bright lighting compared to normal light (Fig. 4D; Supplementary Table S8). These findings emphasize the importance of lighting in automated seed phenotyping using *Samplify*, and we recommend that users carefully control exposure conditions or incorporate suitable controls, such as well-characterized seed populations with predefined abortion ratios. To achieve optimal results, a combination of both approaches is advisable.

Finally, as *Samplify* was mainly developed to provide a quick and unbiased measure to distinguish mutants with different severities of the triploid block, we tested its sensitivity by predicting seed abortion in crosses of a genetic mutant that affects triploid block. Mutants in the MADS-box transcription factor encoding gene *PHERES1* (*PHE1*) and its paralogue *PHERES2* suppress the triploid block. This triploid block is restored when complementing the *phe1phe2* double mutant with *PHE1::PHE1-GFP,* indicating that PHE1 is sufficient to induce the triploid block (Batista et al., 2019). To test whether *Samplify* would identify these differences, we generated the same seed genotypes by pollinating Col-0 maternal plants with pollen from either *omission of second division 1 (osd1)* mutants*, osd1phe2,* or *osd1phe1phe2*. Mutants in *OSD1* produce diploid gametes at high frequency (d’Erfurth et al., 2009) and induce a strong triploid block when used as pollen donor to pollinate diploid maternal plants (Kradolfer et al. 2013). Comparing *Samplify* predictions of F1 Col-0 × Col-0, F1 Col-0 × *osd1*, F1 Col-0 × *osd1phe2,* and F1 Col-0 × *osd1phe1phe2,* we could observe the previously published phenotype of high proportions of fully aborted seeds in F1 Col-0 × *osd1* and F1 Col-0 × *osd1phe2*, while strongly decreased numbers of fully aborted seeds in F1 Col-0 × *osd1phe1phe2* (Fig. 5A; Supplementary Table S9). This significant difference in seed abortion in *osd1phe1phe2* compared to *osd1* and *osd1phe2* was also reflected in the seedling establishment, resulting in nearly 100% of seedlings established in Col-0 and *osd1phe1phe2* pollinations, while only 25.9% and 38.6% in *osd1* or *osd1phe2* pollinations (Fig. 5B; Supplementary Table S10). Taken together, this shows that *Samplify* reliably predicts differences in seed abortion in different genotypes, making it a reliable tool to identify variation in seed abortion between different genotypes.

**Figure 5:**
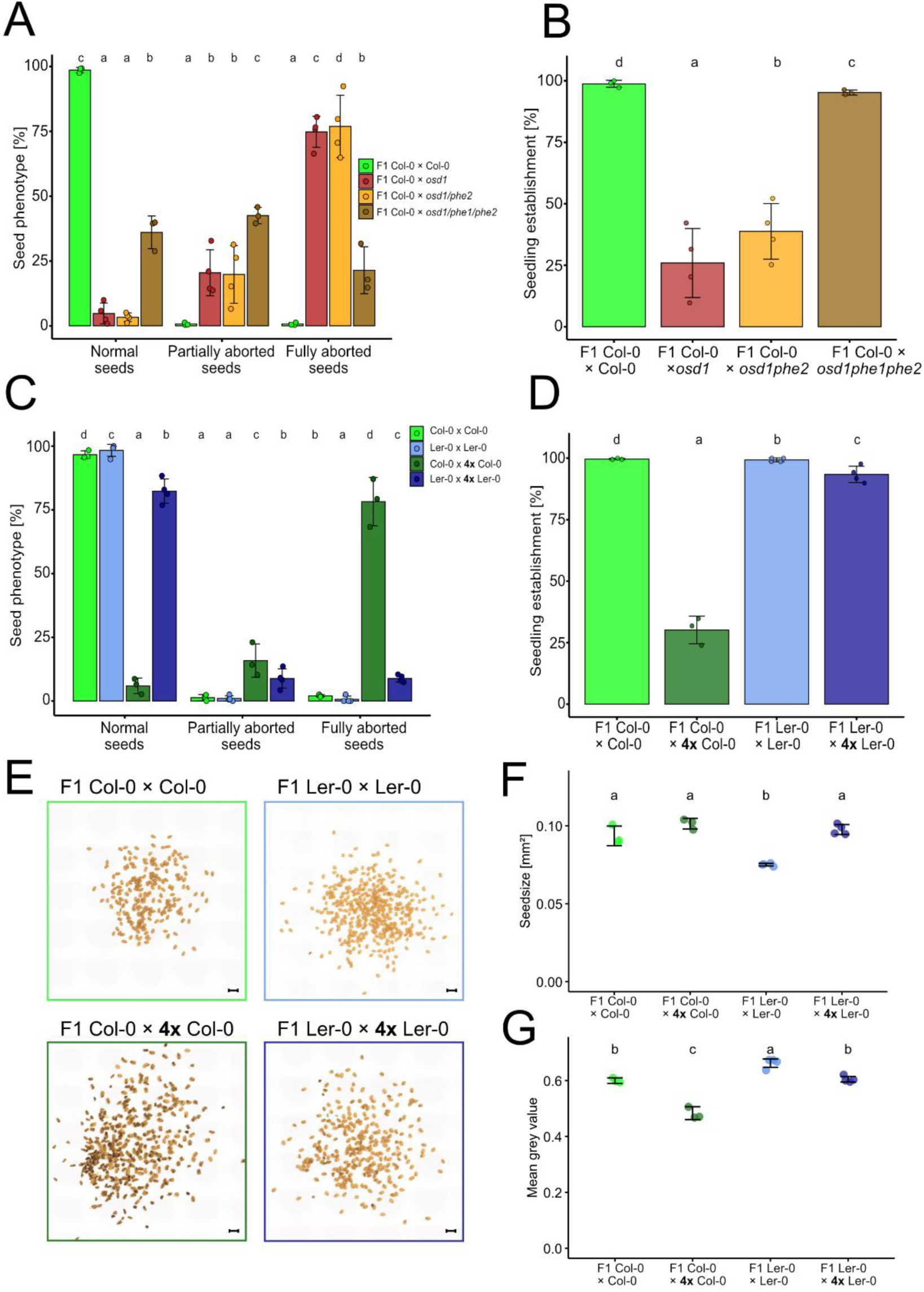
*Samplify* can identify triploid block suppressor mutants and accession-specific differences. A) *Samplify* predicted seed abortion phenotypes in seed populations from different genetic crosses, including relative seedling establishment at four days of growth (B). C) Relative seed abortion phenotype as predicted by *Samplify* in F1 Col-0 × Col-0, F1 Col-0 × 4x Col-0, F1 Ler-0 × Ler-0 and F1 Ler-0 × 4x Ler-0, and seedling establishment at four days of those (D). Statistical comparison was done per category by applying generalized linear models (GLMs) with a Poisson distribution, statistical groups are plotted above (p < 0.05). E) Representative pictures of Col-0 × Col-0, F1 Col-0 × 4x Col-0, F1 Ler-0 × Ler-0 and F1 Ler-0 × 4x Ler-0. Average seed size (F) and average mean grey value (G) of different accession crosses. n = 3-4 pictures. Statistical differences were calculated via one-way ANOVA with post-hoc Tukey test (p < 0.05). Different letters represent different statistical groups. *Samplify* predictions and test statistics can be found in Supplementary Tables S9-S14.

To further test the versatility of *Samplify*, we tested whether it would also reliably predict differences in triploid block between different *Arabidopsis thaliana* accessions. The *Arabidopsis* accession Landsberg *erecta* (L*er*) shows a weaker triploid block compared to the Col-0 accession (Dilkes et al., 2008, Scott et al., 1998). We pollinated L*er*-0 with pollen from autotretraploid L*er*-0 plants (F1 L*er*-0 × 4x L*er*-0) and compared those seeds with their diploid controls (F1 Ler-0 × L*er*-0). Moreover, we also included independent crosses of F1 Col-0 × Col-0 and F1 Col-0 × 4x Col-0, similar to the one shown in Fig. 3. Consistent with previously reported results, we found a lower triploid block in F1 L*er*-0 × 4x L*er*-0 with an average of 81.7% normal seeds, 9.3% partially aborted seeds and 8.9% fully aborted seeds, whereas the F1 Col-0 × 4x Col-0 had 6%, 15.6% and 78.4% of normal, partially aborted and fully aborted seeds, respectively (Fig. 5C; Supplementary Table S11). Also, the average seedling establishment of F1 L*er*-0 × 4x L*er*-0 was significantly higher at 93.4% than that of F1 Col-0 × 4x Col-0 seeds showing only 30.2% of seedling establishment (Fig. 5D; Supplementary Table S12). In conclusion, this data demonstrates that *Samplify* reliably predicts previously published differences of the triploid block between different accessions (Dilkes et al., 2008).

Finally, we tested the application of *Samplify* beyond seed abortion prediction. *Samplify’s* accurate segmentation enables the collection of multiple individual parameters from a given seed picture, facilitating comparisons that do not require prior training of RF models. The L*er* accession is known to have smaller seeds than Col-0 (Herridge et al., 2011, Ren et al., 2019). To test if *Samplify’s* segmentation allows to predict this difference, we extracted the seed area in pixels and converted it to mm^2^. Indeed, this data showed that F1 L*er*-0 × L*er*-0 were significantly smaller compared to F1 Col-0 × Col-0 (Fig. 5F; Supplementary Table S13). Moreover, we utilized the mean grey value from *Samplify’s* parameter list to test the color intensity of F1 seeds. Aborted seeds are usually characterized by a dark brown color, which could be used as a fast assessment of the triploid block, without relying on a pre-trained RF classifier. Indeed, F1 Col-0 × 4x Col-0 had a lower mean grey value compared to F1 Col-0 × Col-0 (Fig. 5G), reflective of high number of aborted seeds in the F1 Col-0 × 4x Col-0. In contrast, the mean grey value of F1 L*er*-0 × L*er*-0 was significantly increased compared to the other F1 populations (Fig. 5G; Supplementary Table S14), consistent with L*er* having a lighter seed color compared to Col-0 (Dilkes et al., 2008). In conclusion, the results presented here show that *Samplify* can reliably identify accession-specific differences in the triploid block, seed size and seed color. The hybrid segmentation of *Samplify* detects the vast majority of seeds in the picture and classifies them based on a pre-trained random forest model with high accuracy, confirmed by germination data as well as published phenotypes. Moreover, since segmentation in *Samplify* is independent of the trained random forest model, key seed parameters such as size or color (grey mean value) can be extracted directly from the parameter list. This enables rapid analysis of different seed populations, making *Samplify* a valuable tool even without a trained random forest model, and extending its utility beyond the sole quantification of aborted seeds.

## Discussion

Plant phenotyping and the (semi-) automated characterization of different traits is a fast-developing area. Quantification of seed phenotypes has been the focus of many research groups, leading to the development of numerous tools over the years (Wang et al., 2023, Wei et al., 2023, Tanabata et al., 2012, Whan et al., 2014). The majority of those tools use simple pictures as input, each usually containing multiple specimens to be described. As such, accurate detection and annotation of all single individuals in a given picture is key. Thus, image segmentation is the first and arguably the most challenging part, it is affected by multiple factors, such as appropriate lighting, sufficient resolution, and often sufficient spacing between single individuals, avoiding touching or overlapping objects. Moreover, the highly irregular shapes of aborted seeds create additional challenges, resulting in frequent misannotation or the loss of substantial numbers.

Here, we present *Samplify* as a tool designed for processing *Arabidopsis* seed images using a hybrid segmentation approach. It combines classical image segmentation techniques with Meta’s Segment Anything Model (SAM). Using this zero-shot learning algorithm, we were able to capture the majority of individual seeds in a single picture, even though they are touching or building larger clusters, which is difficult to identify with classical segmentation approaches. The combination of traditional segmentation together with SAM builds a robust, reliable, and fast tool for seed segmentation, requiring only colored images as input. It leads to high segmentation ratios, aligning almost perfectly with manually counted seeds. On average, *Samplify* processes a picture with approx. 200 seeds in approx.100 seconds, resulting in 0.5 seconds per seed on our system (NVIDIA GeForce RTX 4070 GPU; see material and methods). The Segment Anything Model (SAM) by Meta has been widely applied in medical imaging tasks such as tumor segmentation (Putz et al., 2025) and anatomical structure detection (Lei et al., 2025), but we are aware of only two studies applying SAM in plant phenotyping (Tang et al., 2024, Zhou et al., 2024). Here, we present one of the first applications of SAM for seed segmentation and its first application on *Arabidopsis* seeds.

The combination of balancing classical segmentation approaches and SAM algorithm has been shown to be effective in plant seed phenotyping with the release of GRABSEEDS (Tang et al., 2024). GRABSEEDS, however, was not developed and trained on small *Arabidopsis* seeds and not on seeds in different abortion stages, resulting in morphological irregularities interfering with accurate segmentation. When applied to representative *Arabidopsis* seed images from our collection, GRABSEEDS failed to correctly separate touching seeds. Unlike GRABSEEDS, Samplify does not require or recommend the manual separation of touching seeds, provided they do not lie on top of each other. Seed segmentation is case-specifically balanced between Otsu’s method and SAM. Regions with dense seed clusters are processed using SAM, whereas regions with well-distributed individuals are segmented using Otsu’s thresholding (Otsu, 1979).

*Samplify* was specifically designed for scoring seed abortion occurring in the triploid block, and we provide a trained RF model allowing the quantification of the different seed categories. The model has been trained on about 13000 seeds of different abortion stages in 99 pictures of Col-0 F1 crosses, making it robust for scoring triploid block rates. Using *Samplify*, we were able to predict seed abortion with the same precision as manual counting, which was further validated by germination tests. Moreover, we could show that *Samplify* allows to predict differences in the triploid block of previously identified suppressor mutants (Batista et al., 2019), making it a versatile screening tool for modifiers of the triploid block. Although the RF classifier was trained on Col-0 data, it was able to predict that L*er* has a weaker triploid block(Dilkes et al., 2008). Even though the triploid block detection worked well using our Col-0 based RF model and *Samplify’s* prediction is reliable across multiple biological samples and iterations, we encourage users to train their own models, as seed abortion prediction is model dependent, which in turn, is user and possibly hardware-dependent (imaging).

Segmentation works similarly under different light conditions, yet predictions do vary likely due to changes in the color spectrum, which is, among other parameters, critical for training the RF classifier. *Samplify* corrects for different lighting conditions by applying a global background normalization, nonetheless, well-exposed pictures resulted in the best prediction outcome.

Lastly, it is noteworthy that *Samplify’s* prediction works well for normal or fully-aborted seeds. However, partially-aborted seeds remain the most difficult category to distinguish, as these seeds arise from an ambiguous morphological range, which leads to inconsistencies between human annotations and algorithmic predictions. Partially aborted seeds are the hardest to designate, also for human annotators, as the intermediate phenotype is often misinterpreted as one of the other categories, potentially also reflected in our provided trainings dataset. Nonetheless, *Samplify* reduces the potential human bias, making the predictions more reliable.

*Samplify* was initially developed as a tool for fast and reliable annotation of seed abortion, yet its flexibility and strong segmentation capability makes it useful in many other aspects. For example, we show that *Samplify* can be used for seed size or color comparisons, as exemplified by identifying differences in previously published seed size and color between different *Arabidopsis* accessions (Alonso-Blanco et al., 1999, Dilkes et al., 2008, Herridge et al., 2011, Ren et al., 2019). To facilitate the exploration of different seed shape parameters, we included an extraction function to *Samplify*, enabling users to directly extract specific features used by *Samplify* side-by-side with annotation results.

In conclusion, *Samplify*, together with the pre-trained RF model presented here, offers a fast, accessible, and reliable workflow for quantifying seed abortion across diverse *Arabidopsis* accessions and genetic mutants. By minimizing user-dependent biases and leveraging a hybrid segmentation strategy that achieves near-perfect accuracy while optimizing computational demands, *Samplify* provides a robust, versatile tool for plant developmental studies. Beyond seed abortion, its adaptable framework holds promise for broader applications in seed phenotyping and quantification.

## Material and methods

### Plant material and growth condition

Col-0 and L*er*-0 seeds were obtained from the Nottingham Arabidopsis Stock Center and amplified for multiple generations in the laboratory. Autotetraploid 4x plants were generated via colchicine treatment of seedlings as described (Lafon-Placette et al., 2017). Colchicine treated autotetraploid plants were amplified for at least three subsequent generation before used for crossings. The *phe1phe2* (Batista et al., 2019) in *osd1-3* (Heyman et al., 2011) mutants as well as the *phe2* (SALK_105945; (Batista et al., 2019)) mutant were previously described.

All seeds were surface sterilized with 70% EtOH for 10 min, washed once in fresh 70% EtOH for 10 min, followed by a final wash in 100% EtOH for 5 Min. Hereafter, EtOH was removed by pipetting and seeds were kept in opened tubes under sterile condition until the remaining EtOH evaporated (approx. 30 min). Surface -sterilized seeds were plated on ½ MS plates containing 1% sucrose, 0.68% agar and stratified for two days in the dark at 4°C. Subsequently, seeds were germinated under long-day conditions (16h light/8h darkness) at 21°C and 50 μE light intensity. Seedlings were transferred to soil after seven to ten days and grown in phytotrons under long day conditions (day 21°C, night 20°C 70% humidity, 150 μE light intensity). Crosses were performed about 3-4 weeks after transfer, by emasculating individual flowers, and pollination 48h later with appropriate pollen. Crosses were performed in at least three replicates, whereas each replicate is defined by independent maternal plants pollinated with independent paternal pollen donors. Each cross contained three to five siliques, resulting in about 100-350 seeds. F1 seeds from single crosses were harvested in paper bags when siliques became brown, about three weeks after emasculation. F1 seeds were kept for one week at room temperature before further experiments to ensure proper ripening. Seeds were released from the dry siliques by tapping against the paper bag for a few times. All content was poured on a clean printer pape,r and remaining siliques were removed with forceps. If necessary, seeds were stored in paper bags until imaging.

### Seedling establishment

For the quantification of seedling establishment, F1 seeds from inflorescences containing seeds from 3-5 siliques were evenly distributed on 9 cm round petri-dishes containing 1% plant agar. Hereafter, seeds were stratified for three days at 4°C in the dark. Seedling establishment was imaged after four days of growth (16 h light/8 h dark, 60 µE, 22 °C). Manual counting of seedling establishment was done using ImageJ (Version 1.54f (Schindelin et al., 2012)) using “multi-point tool”, whereas Counter 0 was used for germinating seeds and Counter 1 for non-germinating seeds. Differences in seedling establishment between genotypes were analyzed using a generalized linear model (GLM) with a Poisson error distribution and a log link function.

### Imaging of seed populations

Imaging of seed populations was done using a Keyence Digital Microscope VHX6000 using the ZS20 lens on either 20x Zoom or 50x Zoom. Seeds from single crosses were spread out about 2 cm on clean white paper sheets, making sure that they did not lie on top of each other. Images were acquired in end point stitching mode, scanning an area of about 40 mm × 30 mm to ensure white framing of the seed population. We applied full ring lighting and high exposure to ensure minimal shadows between the seeds. Scale-bars were applied by the Keyence control software and were either 500 um or 1 mm. Pictures were saved in JPEG format. Each experiment was imaged in one session and contained diploid selfed controls and triploid seeds, to ensure the same lighting and edge length across all samples in one experiment. Manual counting for seed abortion was done using ImageJ (Version 1.54f (Schindelin et al., 2012)) using “Multi-point tool”, with Counter 0 for counting normal seed, Counter 1 for the number of partially-aborted seeds and Counter 2 for the number of fully aborted seeds.

To evaluate differences in seed abortion between genotypes, generalized linear models (GLMs) with a Poisson distribution were fitted separately to the counts of normal, partially aborted, and fully aborted seeds using genotype as a fixed effect. For each model, estimated marginal means (EMMs) were calculated using the emmeans R package (Lenth, 2016), which provides model-based adjusted means for each genotype. If applicable pairwise comparisons between genotypes were conducted, and significance groupings were determined using compact letter displays (CLD) from the multcomp R package (Hothorn et al., 2008). Statistical differences in size and color parameters were calculated via one-way ANOVA and post-how Tukey test. All analyses were performed in R (Version 4.5.0;(R Core Team, 2020)), data was processed using the tidyverse R package (Wickham et al., 2019) and plotted via ggplot2 (Wickham, 2009). The multCompView package (Graves et al., 2019) was used for statistical group assignment. All p-values and statistical groups are indicated. Raw values and full test statistics can be found in Supplementary Table S2-S14.

### *Samplify* - Computational Workflow for Seed Segmentation and Classification

The *Samplify* pipeline integrates classical computer vision techniques with transformer-based deep learning models. Images were first converted to grayscale and smoothed using Gaussian blurring (OpenCV 4.7.0)(Bradski and Kaehler, 2008). Initial segmentation applied Otsu’s thresholding(Otsu, 1979) and Canny edge detection (via OpenCV)(Canny, 1986) to separate seeds in sparsely populated regions. For densely clustered seed regions, Meta AI’s Segment Anything Model (SAM, automatic mask generation mode)(Kirillov, 2023) was used, implemented with PyTorch (version 2.0.1)(Paszke et al., 2019) and run with GPU acceleration (CUDA 11.8) (Nickolls et al., 2008). SAM parameters were manually optimized for seed segmentation (e.g., *points_per_side=49*, *pred_iou_thresh=0.86*).

Feature extraction for each segmented seed was performed using a parallelized approach (via Python’s *concurrent.futures.ThreadPoolExecutor*) and included 38 descriptors: 21 color metrics, 9 shape descriptors (e.g., eccentricity, solidity, overlap ratio), 5 Haralick texture features (via *skimage.feature.graycomatrix*), and 3 positional features (Supplementary Table S1).

A Random Forest (RF) classifier (Scikit-learn 1.2.2) (Pedregosa et al., 2011) trained on 13,395 manually labeled seeds from the *TripBlockDefault_RF* dataset was used to classify each seed into one of three categories: normal, partially aborted, or aborted. RF parameters were optimized using grid search, with a test size of 30%, stratified labels, and random state set to 42. The final RF classifier used 100 estimators, maximum depth of 7, minimum samples split of 5, bootstrap enabled, and out-of-bag scoring. Output included color-coded annotated images, feature information for each segmented seed and Excel summary files reporting class counts per image. The pipeline was optimized for high-throughput processing of up to 300 seeds per image and achieved ∼60% reduction in execution time via batch-wise GPU execution and feature parallelization. An overview of the *Samplify* workflow is presented in Supplementary Fig. S1. *Samplify* was tested on two independent systems utilizing either an Intel(R) Xeon(R) CPU E5-2640 v4 @ 2.40 GHz with 20 cores, 64 GB RAM and a NVIDIA GeForce RTX 4070 GPU, or an Intel(R) Xeon(R) W-2133 CPU @ 3.60 GHz with 12 cores, 32 GB RAM and a NVIDIA Quadro RTX 4000 GPU. As *Samplify* occasionally recognized foreign objects, prediction images were manually checked for misannotations. Due to the low number of wrongly annotated objects, we did not apply corrections for full transparency.

## Data availability

*Samplify*, the RF model and detailed user instructions are available on Github: https://github.com/Ronja-Mueller/Samplify.git.

## Acknowledgements

We would like to thank the IT-team at MPIMP for maintaining *Samplify* in the current IT infrastructure. We are grateful for the contributed test and validation data from members of the Köhler group. We acknowledge the assistance of a large language model (LLM) from the GWDG, a high-performance computing center at the University of Göttingen, as well as the use of OpenAI’s ChatGPT (GPT-4, accessed June/July 2025) for assistance with code generation and data processing strategies. All outputs were reviewed and validated by the authors. This work was financially supported by the Max Planck Society.

## Author contributions

Author contributions: H.B., C.K., and D.W. conceived and conceptualized the study; R.L.J.M. wrote and optimized the *Samplify* code and trained the models, A.D. installed and implemented the code base necessary for performing and hosting *Samplify*, H.B. performed the wet lab experiments; C.K., H.B., D.W., R.L.J.M. analyzed the data; H.B and C.K. wrote the manuscript. All authors read and commented on the manuscript.

## Supplementary data

**Supplementary Figure S1:**
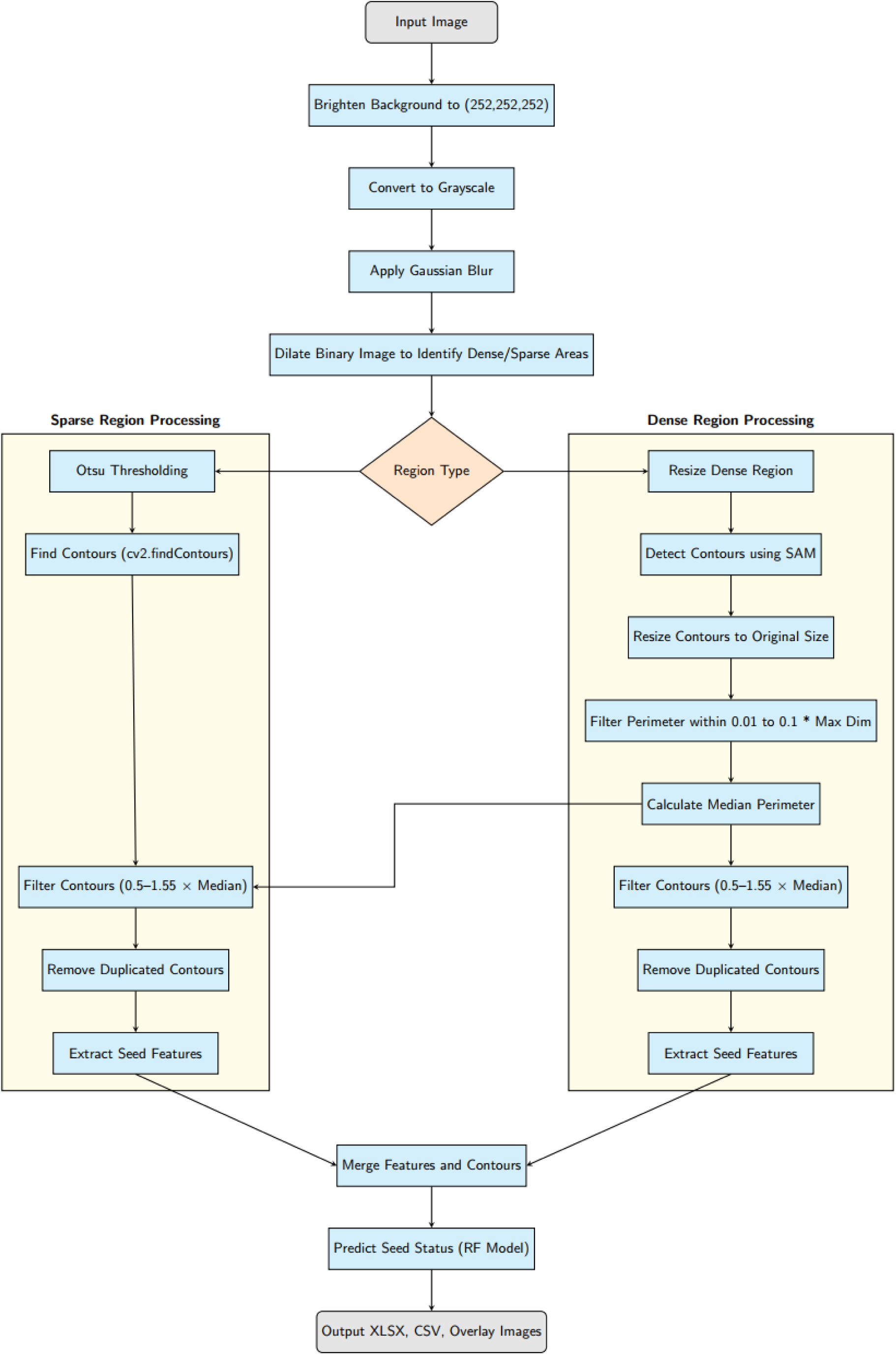
Flowchart of the *Samplify* process.

**Supplementary Figure S2:**
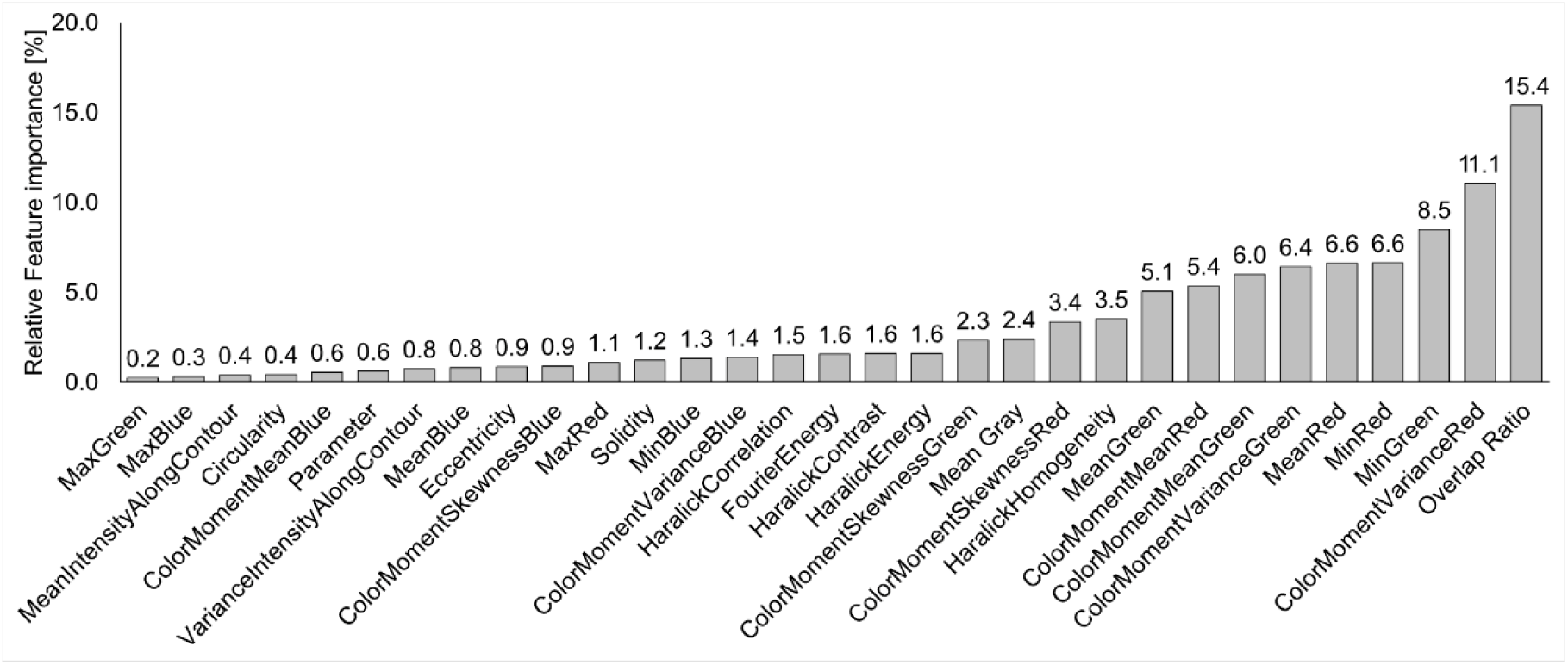
Relative feature importance of all features used by *Samplify* default RF model.

**Supplementary Table S1:** A list of all computed seed shape parameters used by *Samplify* to predict seed abortion.

**Supplementary Table S2**: *Samplify* annotated seeds and test statistics complementary to Fig. 3C. emmean: estimated marginal mean for each genotype; SE: standard error of the estimated marginal mean; df: degrees of freedom for the estimated marginal mean; asymp.LCL: asymptotic lower confidence limit for the estimated marginal mean. asymp.UCL: asymptotic upper confidence limit for the estimated marginal mean. group: statistical group label assigned.

**Supplementary Table S3**: Manually annotated seeds and GLM test statistics, complementary to Fig. 3C. emmean: estimated marginal mean for each genotype; SE: standard error of the estimated marginal mean; df: degrees of freedom for the estimated marginal mean; asymp.LCL: asymptotic lower confidence limit for the estimated marginal mean. asymp.UCL: asymptotic upper confidence limit for the estimated marginal mean. group: statistical group label assigned.

**Supplementary Table S4**: Total seeds counts per picture, counted manually and with *Samplify*.

**Supplementary Table S5**: Seedling establishment results, complementary to Fig. 3F. emmean: estimated marginal mean for each genotype; SE: standard error of the estimated marginal mean; df: degrees of freedom for the estimated marginal mean; asymp.LCL: asymptotic lower confidence limit for the estimated marginal mean. asymp.UCL: asymptotic upper confidence limit for the estimated marginal mean. group: statistical group label assigned.

**Supplementary Table S6**: Seedling establishment results, complementary to Fig. 4A and 4B.

**Supplementary Table S7**: Reproducibility calculation results per seed population and abortion category, complementary to Fig. 4B. SD: Standard deviation; CV: Coefficient of Variation; (SD/mean); CV_percentage: CV*100. Reproducibly: 1-CV. NA indicates instances with less than 10 observations.

**Supplementary Table S8:** Relative *Samplify* prediction results for each category and seed population at different lighting condition, complementary to Fig. 4D.

**Supplementary Table S9:** *Samplify* annotated seeds and test statistics for the triploid block suppressor mutants, complementary to Fig. 5A. emmean: estimated marginal mean for each genotype; SE: standard error of the estimated marginal mean; df: degrees of freedom for the estimated marginal mean; asymp.LCL: asymptotic lower confidence limit for the estimated marginal mean. asymp.UCL: asymptotic upper confidence limit for the estimated marginal mean. group: statistical group label assigned.

**Supplementary Table S10:** Seedling establishment results for triploid block suppressor mutants, complementary to Fig. 5B. emmean: estimated marginal mean for each genotype; SE: standard error of the estimated marginal mean; df: degrees of freedom for the estimated marginal mean; asymp.LCL: asymptotic lower confidence limit for the estimated marginal mean. asymp.UCL: asymptotic upper confidence limit for the estimated marginal mean. group: statistical group label assigned.

**Supplementary Table S11:** *Samplify* annotated seeds and test statistics for the triploid block in different accessions, complementary to Fig. 5C. emmean: estimated marginal mean for each genotype; SE: standard error of the estimated marginal mean; df: degrees of freedom for the estimated marginal mean; asymp.LCL: asymptotic lower confidence limit for the estimated marginal mean. asymp.UCL: asymptotic upper confidence limit for the estimated marginal mean. group: statistical group label assigned.

**Supplementary Table S12:** Seedling establishment results for triploid block in different accessions, complementary to Fig. 5D. emmean: estimated marginal mean for each genotype; SE: standard error of the estimated marginal mean; df: degrees of freedom for the estimated marginal mean; asymp.LCL: asymptotic lower confidence limit for the estimated marginal mean. asymp.UCL: asymptotic upper confidence limit for the estimated marginal mean. group: statistical group label assigned.

**Supplementary Table S13:** Average seed size values in pixel and mm2 as well as ANOVA results of all used accession pictures, complementary to Fig. 5F.

**Supplementary Table S14:** Mean Grey value and ANOVA results of all used accession pictures, complementary to Fig. 5G.

